# Internet of Genetic Things (IoGT): Programmable Long-Distance Cybergenetic Logic Circuits Integrating IoT, Optogenetics, and Bacterial Gene Expression

**DOI:** 10.64898/2026.11.23.768527

**Authors:** Abhishek Paul, Ritwika Basu, Arijit Pal, Biyas Mukherjee, Sangram Bagh

## Abstract

The Internet of Things (IoT) exchanges information and coordinates functions among geographically separated electronic devices. However, establishing analogous long-distance networks between genetically engineered living systems remains a challenge because synthetic genetic intercellular communication largely relies on local molecular signal transport. Here, we introduced a cybergenetic framework named the Internet of Genetic Things (IoGT) that autonomously regulated long-distance gene expression between geographically separated bacterial populations by integrating IoT, bacterial gene expression, and optogenetics. We engineered interconnected IoT nodes containing genetically engineered *Escherichia coli*, in which colour signals arising from reporter gene expression were detected using optical sensors interfaced with a Raspberry Pi. When gene expressions reached predefined thresholds and patterns, data were transmitted over the internet to a remote node, triggering blue-light illumination to induce target gene expression in an optogenetically engineered *E. coli* population. Using this genetic–electronic–genetic communication architecture, we demonstrated cybergenetic BUFFER and OR gates between bacterial populations distributed across nodes separated by 100 m, as well as a cybergenetic AND gate coordinated among three locations. To demonstrate geographically extended operation, we constructed a cybergenetic NOT gate connecting a surface laboratory with a deep-subsurface laboratory located 555 m underground and more than 200 km away. To our knowledge, this was India’s first deep-subsurface biology experiment. IoGT was programmable. It reconfigured the same engineered bacterial populations into diverse Boolean logic gates BUFFER, OR, AND, and NOT gates. These results established IoGT as a programmable cybergenetic architecture capable of remote, multi-node intercellular gene expression control, having significant promise for distributed synthetic biology systems operating across remote or extreme environments.

## 1. Introduction

Bacteria communicate with each other and regulate gene expressions through quorum sensing using chemical signals (Taga and Bassler, 2003). Those chemicals are synthesized and secreted from the bacterial cells, diffused out, and received by neighboring cells to regulate their gene expression in a coordinated manner (Abisado et al., 2018). Such communication governs biofilm formation, virulence, stress responses, and multicellular-like information processing in microbial communities during infections and interkingdom interaction (Rutherford and Bassler, 2012; Whiteley et al., 2017; Abisado et al., 2018; Coolahan and Whalen, 2025).

The advent of synthetic biology (Cameron et al., 2014; “Milestones in synthetic (micro)biology”, 2014) allowed engineering such communication and gene regulation in a human-designed way, including multicellular networks for various programmed pattern formation (Basu et al., 2005; Danino et al., 2010), intercellular toggle switch (Magalhães et al., 2024), building bacterial consortia working as integrated circuits (Perry et al., 2016; Du et al., 2020), and engineered communications between living and artificial cells (Lentini et al., 2017; Smith et al., 2023). Such engineered systems have many projected applications, including treating various diseases (Khan et al., 2022), synthetic microbial communication for bioprocess engineering (Marchand and Collins, 2015; Kang et al., 2022), bioremediation (Li et al., 2021), and engineered communications for soil monitoring in agriculture (Boo et al., 2024). However, those chemical means of cellular communications work only within a short range, like micrometers to sub millimeters distances (Darch et al., 2018; Gestel et al., 2021).

By integrating electronics with engineered biosensor bacteria, scientists developed biodevice to sense gastrointestinal bleeding in pigs by engineered bacteria and responded through luminescence that was detected and wirelessly transmitted by miniaturized electronics (Mimee et al.,2018), a wireless implantable biosensor by integrating genetically engineered *E. coli* with passive microwave backscatter systems (Bilir et al., 2025) to convert molecular sensing into detectable electromagnetic signals in a human body phantom which was received by an external receiver, and engineered bacteria with genetically encoded hyperspectral reporters detected remotely using hyperspectral imaging systems mounted on unmanned aerial vehicles and satellites (Chemla et al., 2025). In those studies, the information change in bacterial behaviors was sent remotely to remote devices for the purpose of human or computer monitoring and did not demonstrate long-distance engineered gene regulations.

Internet of Things (IoT) has emerged as a cyber-physical network of interconnected devices embedded with sensors, processors, communication modules, and software that enable autonomous data acquisition, exchange, and analysis over the internet (Shi et al., 2022; Li and Lao, 2025). For example, in precision agriculture, distributed IoT sensors continuously monitor environmental parameters and plant health, transmitting real-time data to remote servers for analysis and remote monitoring (Mansoor et al., 2025).

Such integration opens new possibilities for constructing wide-area cybergenetic networks in which living bacteria across distinct geographic locations autonomously exchange information at the genetic level and regulate distant gene regulation. Here, by integrating IoT architecture with bacterial gene expression and optogenetics, we introduce the Internet of Genetic Things (IoGT). Using this genetic–electronic–genetic communication framework, we engineered long distance distributed cybergenetic logic gates, including BUFFER, OR, AND, and NOT gates. These enable genetically engineered bacterial populations separated by distances ranging from 100 m to over 200 km (including a deep-subsurface laboratory 555 m underground) to autonomously communicate over the internet and regulate gene expression in remote populations.

## 2. Results and discussion

### 2.1 The IoGT Architecture and Genetic Systems Design

Our IoGT (Fig. 1) was designed using geographically separated IoT nodes, each consisting of a Raspberry Pi 4B. The sender nodes (node Xi) interface with populations of genetically engineered *E. coli* cells on petri dishes that produce a colored protein as a function of a chosen chemical inducer. In node Xi, a small (22 mm X 22 mm) optical RGB colour sensor is positioned above the petri dish using a custom built, height adjustable sensor holder (Fig. S1 a, b).

**Fig 1.**
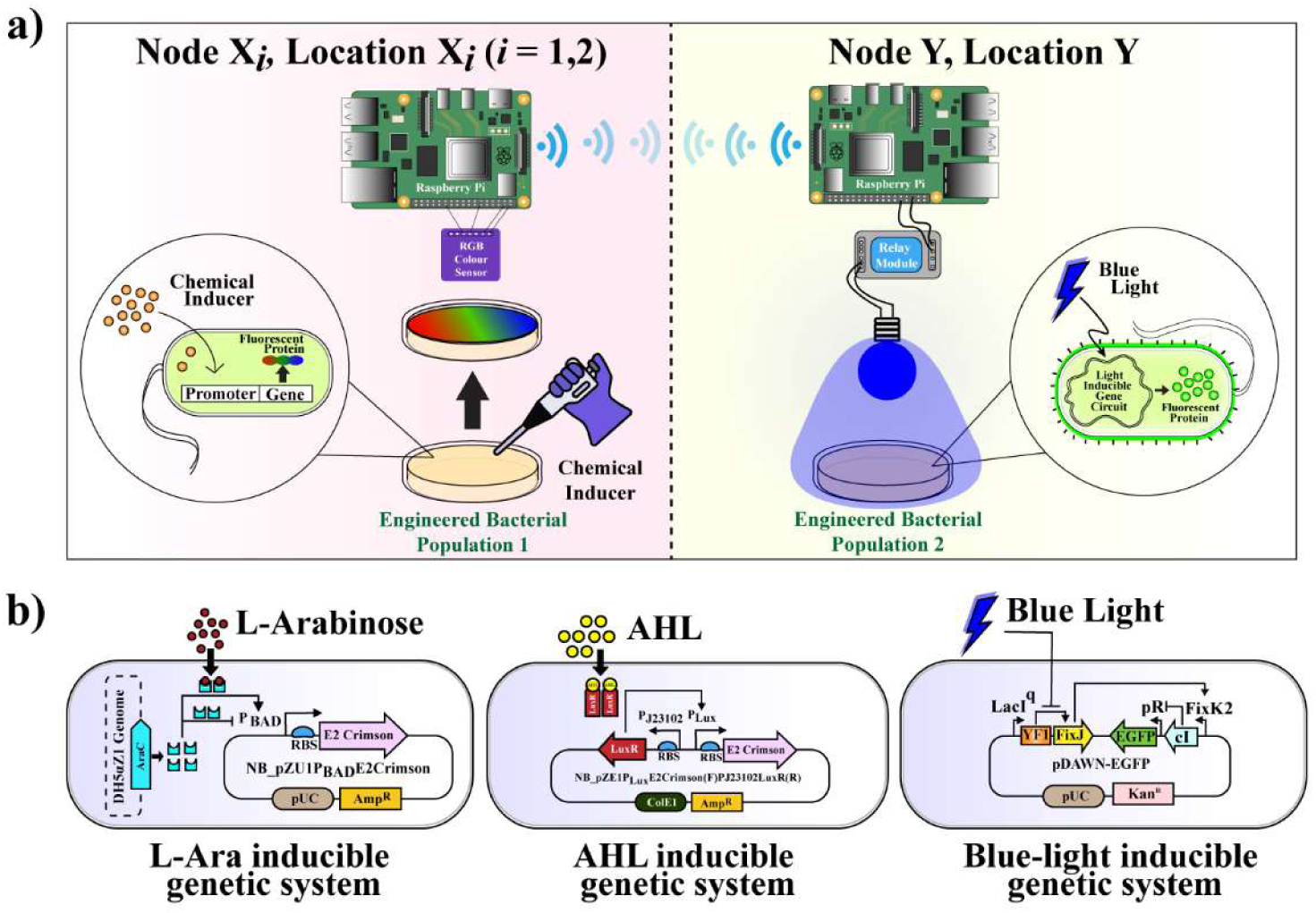
**The schematic design of loGT**. a) IoT is integrated with bacterial gene expression through optogenetics. At node Xi, a chemical inducer is added to a lawn of engineered bacterial populations. Expression of a fluorescent protein resulted in a colored bacterial lawn. The color change is detected and collected by an RGB color sensor attached to a Raspberry Pi placed directly above the lawn. The information is transmitted via the internet to a second location, where a blue LED light is turned on automatically through a relay switch and Raspberry Pi. An optogenetically engineered bacterial population at this location gets induced via blue light and expresses a target protein. B) Gene systems design of L-arabinose, AHL, and blue light inducible cells.

The sensor is wired to a Raspberry Pi 4B. Custom Python scripts running on the Raspberry Pi continuously acquired red, green, and blue (RGB) values through I2C communication and calculated mean color intensities from time-resolved measurements. The RGB sensor continuously captures color information as an optical output signal whose intensity should reflect the internal gene-expression state of the induced *E.coli* population. When the optical output crosses a predefined threshold and a pattern (such as above-threshold expressions from two locations), the system automatically transmits a trigger message over the internet to the receiving IoT node (node Y). Node Y, consisting of another Raspberry Pi, receives and interprets the incoming signal and uses it to activate a solid-state relay, which controls an external blue LED light source (Fig. S1c,d). The parts and the assembled nodes Xi and Y are shown in figure S1. The details of the configuration, coding, and assembly for node Xi and node Y are discussed in the methods section. The blue LED in the location of node Y serves as an optical stimulus to an engineered *E. coli* cell population containing a blue light-responsive genetic circuit. The optogenetically engineered bacteria would process that transferred information and respond to it by altering a target gene expression. After assembling and configuring various parts, we tested and optimized the node Xi for RGB sensor data capturing with various colored papers (Table S1). The optimum distance from the colored object and sensor was found to be ∼ 10 mm. The wireless communication between node Xi and node Y was tested with colored papers, varying the thresholds for blue values (20, 40, 60, 80, 100). The minimum lag time between sensing blue at node Xi and turning on the blue light transilluminator at node Y remotely and automatically was found to be 15 seconds (Video S1).

Next, we engineered three genetic systems; an arabinose inducible, an AHL inducible and an blue light inducible gene expression system (Fig. 1b). In one population, *E. coli* DH5αZ1 cells (Lutz and Bujard, 1997) expressed fluorescent protein E2-crimson (Strack et al., 2009) under arabinose induction. Here, E2-crimson gene was placed under the control of arabinose induced P_BAD_ promoter in a plasmid with a pUC origin of replication. The plasmid was inserted in *E. coli* DH5αZ1 cells. The AraC protein required for the induction of pBAD with arabinose was inherently supplied by the DH5αZ1 genome. In another system *E. coli* DH5αZ1 expresses E2-crimson under AHL induction. Here, the E2-crimson gene was placed under a pLux promoter. A pLux promoter in the presence of the LuxR protein and AHL molecule gets induced. The LuxR gene was placed under a constitutive J_23102_ promoter in the same plasmid with ColE1 Ori and transferred to *E. coli* DH5αZ1. The blue light induced bacteria (Fig. 2b) were engineered based on the pDAWN optogenetic system (Ohlendorf et al., 2012) consisting of a YF1-FixJ operon, where, in the dark, the histidine kinase YF1 phosphorylates FixJ, which represses downstream expression by binding with the promoter pFixK2, and blue light inhibits YF1 from phosphorylating due to the change of conformation of YF1. We placed an enhanced green fluorescent protein (EGFP) gene as a target gene under a pR promoter, which could be repressed by the lambda repressor protein CI. The CI protein was under pFixK2, resulting in expression of EGFP under blue light. Both the chemically inducible cells would work as sender cells, from which the information of E2-crimson expression would be transmitted and the blue light inducible system work as the receiver cells, where the target gene would be expressed based on the gene expression thresholds and patterns from the sender cells.

**Fig. 2.**
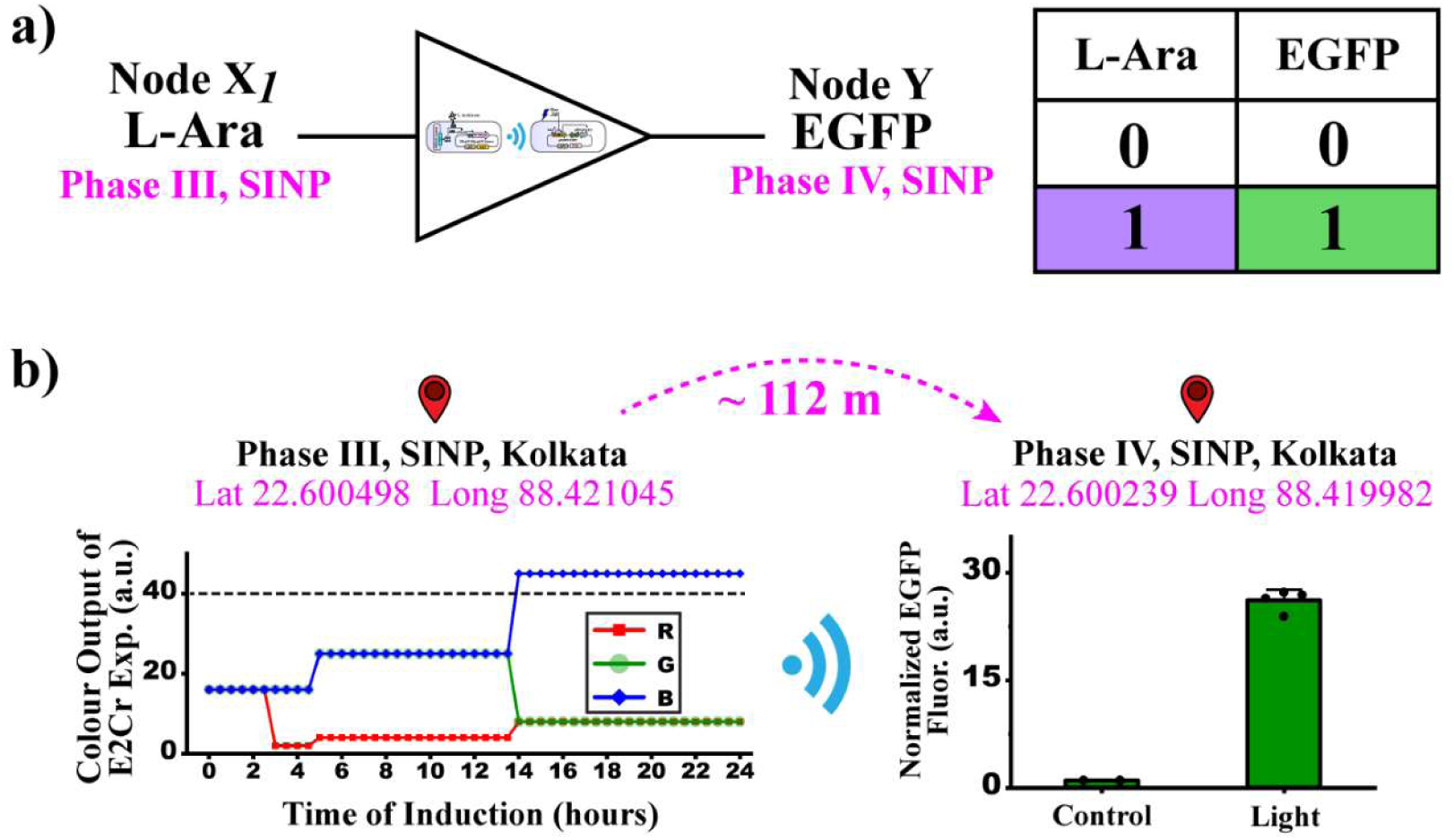
Long distance intercellular BUFFER gate. . a) Biochemical BUFFER gate logic and truth table. b) RGB profile of L-arabinose induced E2-crimson expression (biological replicate I) at Node Xi in phase III, SINPc) Target EGFP expression from optogenetically engineered bacterial cells after 6 hours of blue light induction at a Phase IV lab at SINP, 112 meters apart.

### 2.2 Long-distance intercellular BUFFER gate with genetic parts separated by a hundred meters apart

With this system architecture and genetic systems, first, we built a long distance intercellular BUFFER gate (Fig 2). We placed two engineered bacterial populations along with the two IoT nodes in two different physical locations with an aerial distance of 112 meters (Fig. S2).

Engineered bacteria from a single colony were grown overnight, diluted 100X, and resuspended in 100 mL LB media with appropriate antibiotics and grown for ∼ 5 hours (OD_600_ ∼ 0.4-0.5). Then the cells were plated on LB agar medium with antibiotics in a petri dish and grown at 37^0^ C. After overnight growth of a bacterial lawn, the plate was placed in location X1 (Phase III, SINP) with node X1 inside an incubator (37^0^ C), and 2mL of L-arabinose (10% w/v) was added to the lawn. A wet chamber with ddH_2_0 was placed nearby to maintain the humidity. Petri plate lids were modified to have a square aperture (15 mm X 15 mm) in their center to allow the RGB sensor to collect colored gene expression information from the lawn surface. The RGB sensor on top of the petri dish was connected to a Raspberry Pi and continuously read all three color outputs (red, green, and blue) from the engineered bacterial culture (Fig. 2b). We recorded the measurements every 30 minutes, an average of 10 seconds in digitized format. The expression of E2-Crimson increased with time (Fig. S2b) and was continuously measured with the RGB sensor and Raspberry Pi (Fig. 2b). We set an arbitrary threshold value of 40 for the color blue; when reached, it automatically transmitted a signal to the other location (location Y, node Y) through the internet (Fig. 2b, Fig. S2a). The transmitted signal was received on a second physical location (Phase IV, SINP) by the remote Raspberry Pi based IoT node Y and routed to a solid-state relay acting as a digital actuator (Fig. S1d). Upon activation, the relay switched on a blue LED transilluminator positioned inside a controlled-environment incubator (37^0^ C, ∼ 60 RPM). Four biological replicates of optogenetically engineered *E. coli* (Fig. 1b) were grown separately overnight in LB medium with antibiotics, diluted 200X, and suspended in 20 ml of fresh M9 liquid medium with antibiotics in four petri dishes and were placed on the blue LED light transilluminator before adding the arabinose at the bacterial population in location X1. The LED provided the defined optical stimulus for the second bacterial population.. After the cells were placed in both locations and the cells at the first locations were induced, no human intervention was done till the experiment was complete.

We observed that the blue color threshold for E2crimson gene expression in arabinose-induced E. coli cells was crossed around 14 hours after induction (Fig. 2b). As the engineered bacterial cell population at location 1 crossed the threshold value (40), the system automatically sent a signal over the internet, and the blue light was automatically turned on at location Y. A representative E2Crimson expression profile from a lawn of arabinose-induced bacteria is shown in Figure S2b. The cell plates on the blue-light transilluminator were collected after 6 hours, spun down, and washed twice with phosphate-buffered saline (pH = 7.3). EGFP expression in the cells was measured using a fluorescence microplate reader and showed a 26-fold change between these light-induced cells and control cells placed in the dark (Fig. 2b). The control experiment with engineered cell population 1, without arabinose induction, showed that the RGB color value remained at the basal level under the same experimental conditions (Fig. S2a). The long-distance intercellular gene regulation experiments and their controls were performed with two additional biological replicates (three in total) on different days, each yielding similar observations (Fig. S3). This demonstrated that long-distance intercellular communication and gene regulation can occur between two bacterial populations placed approximately one hundred meters apart, with a change in gene expression at one location influencing gene expression in another population 112 meters away autonomously.

### 2.3 Long distance intercellular OR gate with genetic parts separated by a hundred meters apart

Next, we showed that the same system architecture can be used to build a long-distance intercellular synthetic genetic OR gate, such that three genetic parts distributed among three engineered cell populations placed in two different physical locations (Fig. 3a). The combinations of two input chemicals would be applied to the engineered ‘input cells’ at one physical location, and the logical output of reporter gene expression would be obtained from the ‘output cells’, placed in another physical location 112 meters apart (Fig. 3a).

**Fig. 3.**
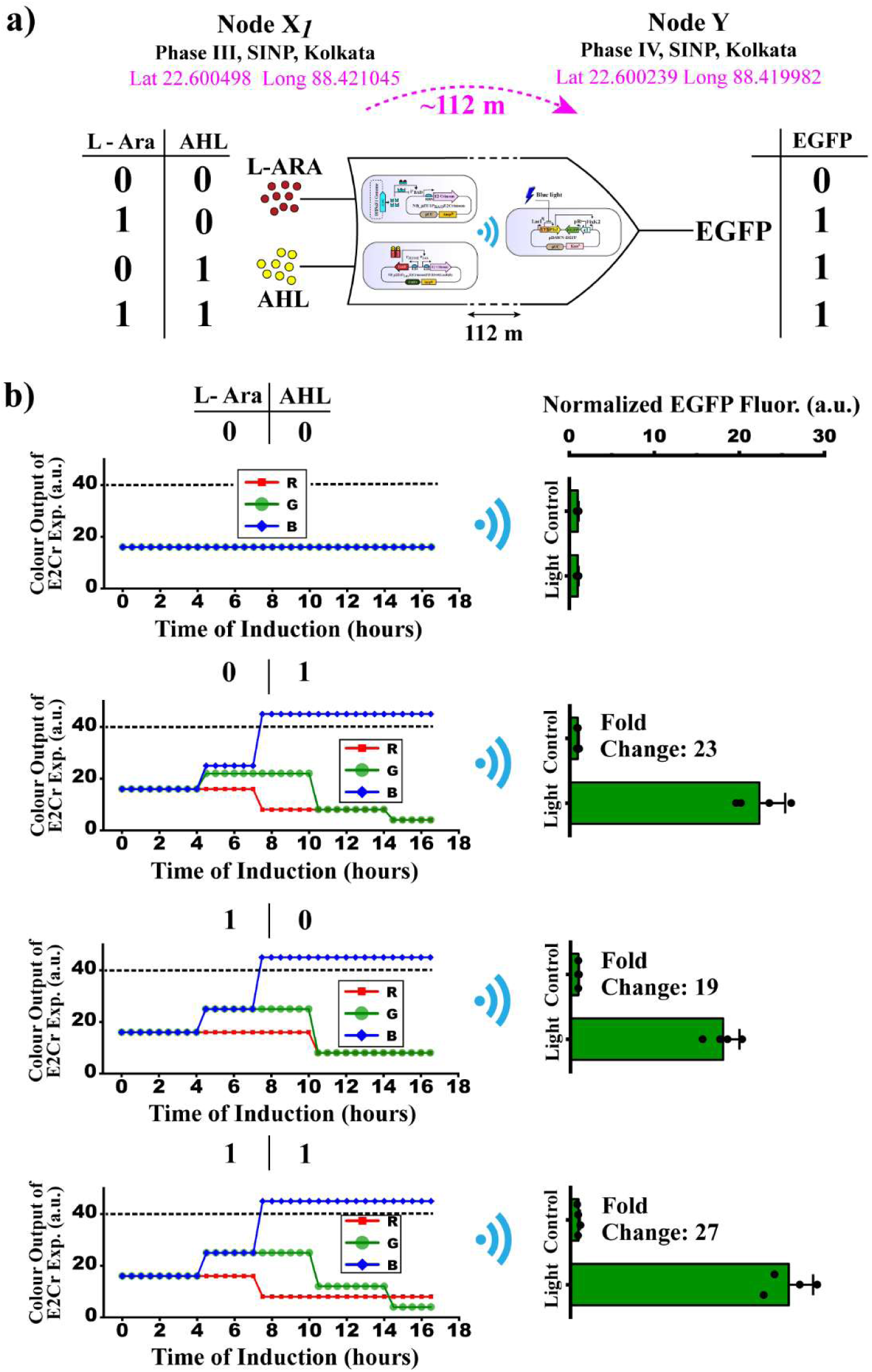
Long distance intercellular OR gate. a) Schematics of the long distance intercellular OR gate. The input chemicals will be applied in location 1 to a mixed population of L-arabinose and AHL induced cells. The logical output will be obtained from location 2 by EGFP expression. The truth table for the OR gate is shown. b) Experimental results of the long-distance OR gate. The E2Crimson expression profiles at location 1 and EGFP expression profiles at location 2 are shown as a function of various combinations of input chemicals.

For OR gate, apart from the arabinose induced cell, we used another engineered set of *E. coli* cells, which produced E2-crimson upon AHL induction (Fig. 1b, S4a). To test the AHL incuced cells, we grew a lawn of AHL induced engineered cells on an LB agar plate from a single colony and induced it by adding 2 mL of AHL (125 µM) to the lawn. Next, we checked the E2-crimson expression from those induced cells with the same RGB sensor system by measuring the red, green, and blue color values continuously (Fig. S4b). Then we checked how the initial number of cells for building the bacterial lawn influenced the time to reach the threshold. We observed that increasing the number of cells approximately 2.5 times would reduce the time to reach the threshold around half for arabinose induced cells (Fig. S4 c, d). Therefore, we decided to use the increased cell numbers for the OR gate experiment to reduce the threshold time around 7 hours.

Next, we grew a mixed population of both arabinose and AHL induced cells on an LB agar plate. This mixed cell population at node X1 (location X1) served as the input cells, and the optogenetically engineered light induced cells at node Y (location Y) worked as output cells. We placed the input cells under node X1 and added appropriate inducer combinations (L-Ara = 0, AHL= 0; L-Ara = 0, AHL=1; L-Ara =1, AHL= 0; L-Ara =1, AHL=1) (Fig. 3b). Here, inducer condition ‘0’ means absence of a specific inducer, and ‘1’ means higher than saturated concentration (10 % w/v for L-arabinose and 125 µM for AHL).

Keeping the threshold value at 40, we observed that the output cells at location 2 showed a high EGFP expression (the ON state) when either AHL or arabinose or both were added to the input cells at location 1 (Fig. 3b). A zero or low basal level of expression (the OFF state) was observed in the cells at location 2 when both inputs were absent in location 1. The results suggested the successful long-distance synthetic genetic OR gate behaviour following the truth table of a logical OR between AHL and arabinose.

### 2.3 Long distance synthetic genetic AND gate: Intercellular gene regulation between three locations

Next, we demonstrated a long distance synthetic genetic AND gate. Three genetic parts were distributed among three engineered cell populations and were placed in three different physical locations under three IoT nodes, namely node X1, X2, and Y (Fig. 4a). We used the same three engineered cells used for OR gate (Fig 1b, Fig 3a). We placed the arabinose induced cells under node X1, AHL induced cells under node X2 and the optogentically engineered output cells under node Y. Arabinose and AHL induced ‘input cells’ were separated by around 40 meters, and the output cells were placed around 100 meters away from the both input cells. (Fig. 4a).

**Fig. 4.**
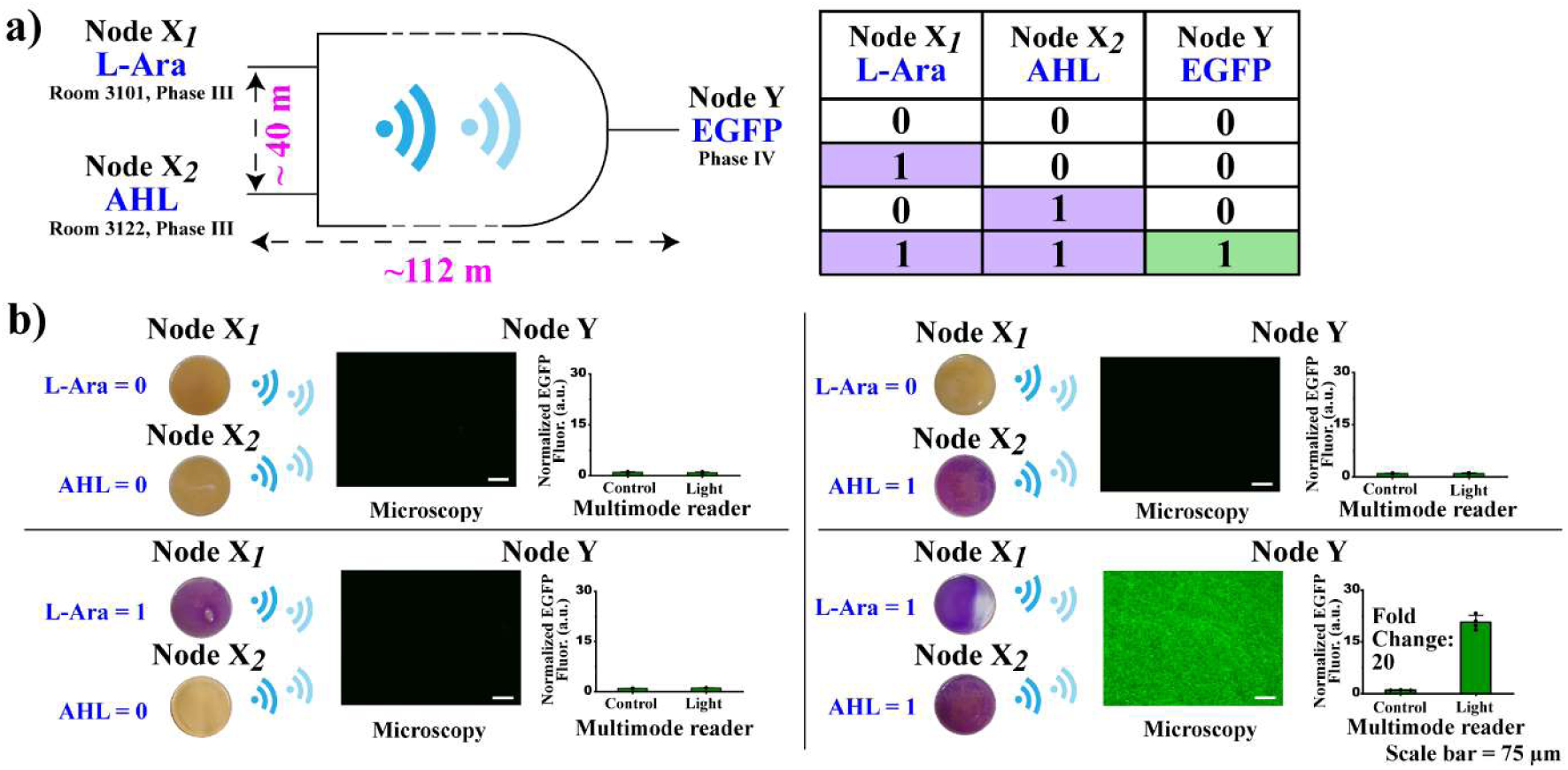
**Long distance intercellular AND gate**. a) Intercellular AND gate logic and truth table. b) Experimental results of the long-distance AND gate. The output EGFP expression profiles as a function of various combinations of input chemicals L-Ara and AHL. The coloured bacterial lawn showed various expression patterns of E2-crimson. The logical output in the form of EGFP expression is shown through fluorescence microscopy images of the bacterial population and the bar diagram beside it shows the mean of quantitative measurements of EGFP expression from four biological replicates using a multimode plate reader from the same experiment.

We updated the computational rules at the IoT layer for AND gate such that when above threshold gene expressions data transferred from both X1 and X2 locations after a predefined time (12 hours) only then the relay switch turned ON the blue light. Input chemicals were added appropriate locations and combinations (L-Ara = 0, AHL= 0; L-Ara = 0, AHL=1; L-Ara =1, AHL= 0; L-Ara =1, AHL=1) (Fig. 4b). Keeping the blue threshold value at 40 for E2-crimson expression, we observed by multimode reader and with epifluorescence microscopy that the output cells at location Y showed a high EGFP expression (the ON state) only when both inducers were added to the input cells at location X1 and location Y (Fig. 4b,c). A zero or low basal level of expression (the OFF state) was observed in the output cells at location Y when either input was absent in distant locations. The colour (RGB) profile, DIC images and AND gate experiments with biological replicates have shown in figure S5. The results suggested the successful IoT enabled long-distance synthetic genetic AND gate behavior, where target gene regulation of a cell population in a specific location was controlled by gene expression pattern of two different cell populations in two other physical locations. This suggested that unlike purely chemical based intercellular synthetic genetic logic gates and devices (Tamsir et al., 2011, Ausländer et al., 2018, Padmakumar et al., 2025) that required distinct genetic constructs for different computational operations like OR and AND gate, in our system we could use the exact same engineered bacterial populations into different Boolean logic gate like OR and AND gate by updating the computational rules at the IoT layer.

### 2.4 Long-distance genetic NOT gate between a deep subsurface underground lab and a surface lab in two cities

Deep subsurface underground laboratories are unique research facilities that provide a lower background radiation environment than the Earth’s surface. Biological experiments performed in deep underground facilities such as Gran Sasso (Italy) and Baksan (Russia) have revealed intriguing phenomena, demonstrating that ultra-low radiation background (ULRB) conditions reduce cellular protection against DNA-damaging agents in yeast and mammalian cells, while triggering distinct stress responses in Drosophila and C. elegans (Satta et al., 1995, Esposito et al., 2020, Castillo and Smith, 2017, Zarubin et al., 2021a, Zarubin et al., 2021b). Because manual intervention in such extreme, isolated environments is difficult, our IoGT system provides an ideal platform to perform such biology experiments remotely and autonomously.

To test the scalability of our system over increasing distances, we first performed experiments between two surface laboratories separated by an aerial distance of ∼212 km: the Jaduguda Health Physics Laboratory, Jaduguda and the Saha Institute of Nuclear Physics (SINP) in Kolkata, demonstrating successful long-distance communication (Figures S6, S7, and S8).

Building upon this long distance connectivity, we next demonstrated a distributed cybergenetic NOT gate connecting the deep subsurface underground lab (JUSL) at Jaduguda (located 555 m underground) with the surface laboratory at SINP, Kolkata (Figure 5). JUSL is a physics facility dedicated to dark matter searches (Das et al., 2025). For this experiment, AHL induced sender cells were placed inside an incubator within the deep-subsurface facility under node X1, while optogenetically engineered receiver cells were maintained under node Y in the SINP, Kolkata surface laboratory, with remote communication established via VPN. When AHL was omitted from the deep-subsurface culture at JUSL, cells at the SINP surface lab expressed EGFP, and vice versa. The RGB profile, DIC and merged microscopy images, and results with biological replicates have shown in Figure S9. This operational loop successfully establishes a functional cybergenetic NOT gate operating between a deep-subsurface environment (555 m underground) and a remote surface laboratory separated by over 200 km. To the best of our knowledge, this was the India’s first deep-subsurface biology experiment.

**Fig. 5.**
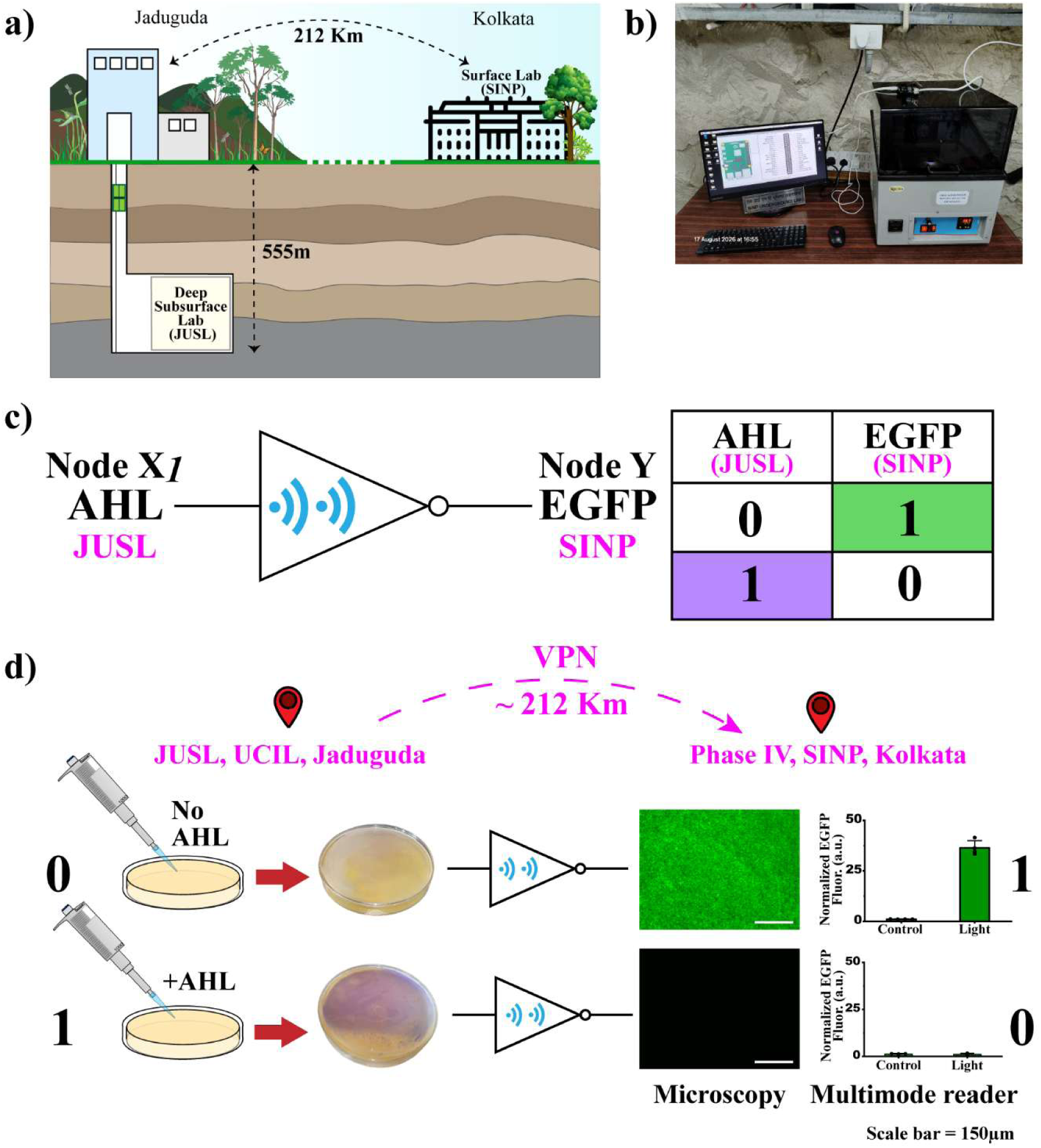
a) A schematic of JUSL underground and SINP surface lab. b) Experimental setup at JUSL. c)Intercellular NOT gate logic and truth table. d) Experimental results of the long-distance NOT gate between 555 m deep underground subsurface lab and surface lab 200 km apart. The output EGFP expression profiles as a function of AHL induction. The coloured bacterial lawn showed expression patterns of E2-crimson. The fluorescence microscopy images of the bacterial population shows the logical output in the form of EGFP expression and the bar diagram beside it shows the quantitative mean EGFP expression from four biological replicates using a multimode plate reader from the same experiment.

## 3. Conclusions

Taken together, we have demonstrated the Internet of Genetic Things (IoGT), an IoT-enabled cybergenetic framework that achieves autonomous, long-distance intercellular gene regulation across geographically separated bacterial populations. By interfacing optical detection of chromogenic reporter gene expression at sender IoT nodes with automated thresholding, internet transmission, and optogenetic blue-light actuation at remote receiver nodes, we converted cellular gene expression events into digital signals that directly govern target gene expression in distant living cells. Using this genetic– electronic–genetic communication architecture, we implemented distributed cybergenetic BUFFER and OR gates across nodes separated by 100 m, a three-location cybergenetic AND gate, and a long-distance cybergenetic NOT gate connecting a surface laboratory with a deep-subsurface facility located 555 m underground and more than 200 km away. To our knowledge, this was India’s first deep-subsurface biology experiment. Importantly, because logic rules are programmed at the IoT layer, the same engineered bacterial strains could be repurposed across multiple distinct logic gates without modifying their underlying genetic circuitry.

This IoGT capability is expected to become increasingly vital for biological explorations in hard-to-reach locations and extreme environments like space habitats (Elsaesser et al., 2023), deep-subsurface laboratories (Esposito et al., 2020), and ocean floor nodes (Jones et al., 2025). Furthermore, it may address critical industrial and ecological challenges by enabling autonomous, coordinated microbial behaviour for wide scale environmental sensing, real time bioremediation, adaptive bioprocess control, and distributed biological computation(Sarkar et al., 2021; Canadell et al., 2022; Bonnerjee et al., 2024; Padmakumar et al., 2024) across geographically isolated nodes. Ultimately, this work establishes wide-area cybergenetic networks operating at the gene regulatory level, having strong potential for building next generation distributed synthetic biology systems with many applications.

## 4. Materials and methods

### 4.1 Genetic Constructs

Standard molecular biology protocols were used to build all the genetic constructs. The genetic constructs were built using the Network Brick (NB) assembly principle with NB plasmids (Mukhopadhyay et al., 2020). The arabinose induced construct was built in a pUC19 NB plasmid, pNB_EA_3EGFP^37^. The promoter P_BAD_ and E2-crimson gene were placed between XhoI and EcoRI and between KpnI and XbaI sites, respectively. The AHL induced construct was built by placing two genetic cassettes in a single plasmid in opposite directions in an NB plasmid. In one cassette, the E2-crimson gene was placed under a P_Lux_ promoter, and in the other cassette, a LuxR gene was placed under a constitutive promoter, P_J23102_. The optogenetic construct was built using the pDawn system (Ohlendorf et al., 2012), wherein in a pDawn empty vector (Addgene plasmid # 43796), EGFP was placed under the P_R_ promoter. The plasmid details are tabulated in table S2. KOD Hot-start DNA Polymerase (Merck) was used for all PCR reactions. All restriction enzymes, buffers, ligase, and ladders used in the work were purchased from New England BioLabs. Plasmid isolation, PCR purification, and gel extraction kits were from Qiagen. The primers were constructed from IDT. The constructs were sequence verified from Eurofins Genomics India Pvt. Ltd. The details of the plasmids, bioparts and oligos used in this study are tabulated in Table S2, S3, and S4.

### 4.2 Chassis Organism and Bacterial Culture

All clonings were performed in the *Escherichia coli* DH5α strain. All experiments were performed by transforming *E. coli* DH5αZ1 strain cells with the constructed plasmids. For initial fluorescent gene expression experiments in liquid culture (Fig. S7), four colonies for each construct were grown overnight in Luria-Bertani medium (LB), Miller (Himedia) with 100 μg/mL ampicillin /50 μg/mL kanamycin (Himedia) as appropriate at 37 °C, with 250 RPM shaking, diluted 100X in fresh LB liquid media with appropriate antibiotics and inducers (1% w/v L-arabinose Himedia / 5 µM AHL C6-HSL, Sigma-Aldrich / 6 hours blue light from a LED based blue light transilluminator, Invitrogen) and regrown for 16 hours (arabinose and AHL induced cells), and regrown in 20 mL of M9 media (blue light inducable cells) for 6 hours in 90 mm perti dishes with 60 RPM shaking for light induced construct (Fig. S7). For long distance communication experiments, overnight cultures are diluted 100X and grown in 100 mL or 250 mL media (as mentioned in the main text) and regrown for ∼ 5 hours (OD_600_ = ∼ 0.4-0.5) and then plated on freshly prepared LB-agar plates with antibiotics. After growing the lawns overnight at 37 °C, appropriate inducers were added (2 mL of 10% w/v L-arabinose/ 125 µM AHL). The time of adding the inducer is the time ‘0’ for the long-distance communication experiments. For OR gate experiments, separate overnight cultures (AHL induced and arabinose induced) were diluted and regrown in 125 mL LB separately, and after washing, both the cells were spread on the LB-agar plate together.

### 4.3 Fluorescence Characterization in Multi-Mode Plate Reader

Fluorescence outputs from the reporter proteins for liquid culture experiments were measured in Synergy HTX Multi-Mode reader (Biotek Instruments, USA). After the mentioned period of induction for the three constructs, cells were spun down, washed, and resuspended in 1x PBS (phosphate buffered saline; pH ∼ 7.3) and loaded into a black flat-bottom 96-well plate (Greiner/Falcon) and OD_600_ value and fluorescence were measured using appropriate combinations of filters for E2 Crimson (610/10 nm band-pass excitation and 645/10 nm bandpass emission filters) and EGFP (485/20 nm excitation and 516/20 nm emission filters) with gain kept at 70 for both the reporter genes. For estimating autofluorescence background, E. *coli* DH5αZ1 cells without any plasmids were measured.

### 4.4 Fluorescence Data Analysis

For determination of fluorescent gene expression, the absolute value (a.u.) of fluorescence for each bacterial culture was normalized to the number of cells by dividing it by its respective OD_600_ value:

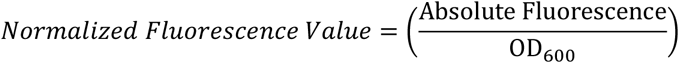

Next, mean autofluorescence from DH5αZ1 cells carrying no plasmids was subtracted from the normalized fluorescence value of individual cells of the experimental sets. Mean normalized fluorescence intensity without autofluorescence was finally calculated for each experimental set by:

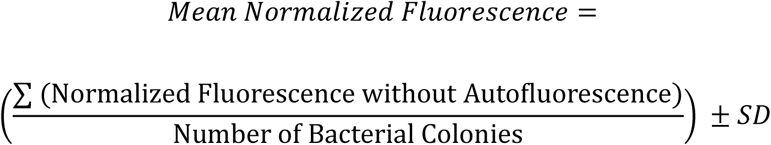

Fold change of gene expression was measured by:

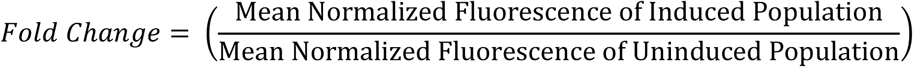

All data analysis and graphs were prepared using OriginPro 2026 (OriginLab Corporation, USA) software.

### 4.5 RGB Color Detection and Transmission

To build node X*_i_* for the detection of color resulting from E2 Crimson expression from the engineered bacterial lawn, an RGB color sensor, TCS34725, was used. The sensor was integrated with a Raspberry Pi 4B (RPi) via jumper cables. To enable the connection between the two, the RPi was configured to enable an I2C connection. Once the I2C connection was enabled, libraries like RPI.GPIO, which provides Python code to utilize the GPIO pins on the RPi, and the adafruit-circuitpython-tcs34725 library, which provides packages containing Python codes to interface the sensor with the RPi and retrieve color data, were installed in the RPi. After installing the necessary libraries, Python code was written that creates a.txt file containing RGB values sampled each second for ten seconds with a datetime stamp and gives a mean RGB value of the 10 readings every time the code is executed. This code was made to execute once every 30 minutes via the Cron task scheduler in the Linux based RPi. The color information as and when received was stored in Dropbox. Another set of Python codes created flag files and recorded values if the predefined threshold of E2crimson expression was reached from a specific node.

### 4.6 Receiving Node and Relay Switch

The receiving end node (node Y) contained another Raspberry Pi in combination with a relay switch. This Raspberry Pi scanned for the files in the dropbox, which was created in the node Xi and transferred them over the internet. To control the switching ON/OFF of a blue light source, a Raspberry Pi in combination with a relay switch has been used. Since the GPIO pins of an RPi operate at low voltages and cannot directly drive high-powered electrical devices like an LED light source, a relay switch acts as an electrically isolated switch between the two. The relay switch was wired to the RPi and the light source via jumper cables. The connection was established by installing necessary libraries. To configure the BUFFER, OR and AND gates, a python code was written which, when executed, scanned Dropbox for the flag file(s) created by Node X*_i_*. On detection of the file(s) in Dropbox, and following the respective Boolean logics, the relay was switched ON, which in turn switched ON the blue-light source. For cybergenetic NOT gate between deep subsurface lab and surface lab, instead of a dropbox the local desktop was scanned *via SSH.* The codes were executed via Cron every minute for BUFFER and OR gates and after 12.5 hours for AND and NOT gates after adding inducers at Node X*_i_*. Accessory addons like display monitor, keyboard and mouse, Ethernet and HDMI cables were also connected to the Raspberry Pis used in both nodes. The codes are available in GitHub (https://github.com/syntheticbiologylabsinp/Synthetic-long-distance-cell-cell-communication).

### 4.7 File creation and detection in Dropbox

To create files and scan for them in Dropbox via Python programming, a Dropbox application was created via the App Console menu of the Dropbox Developer website. Necessary permissions and settings were configured that allowed the Python codes to read and write files in the Dropbox. A refresh token was generated using the app key, app secret and access token and included in all the codes for continuous uninterrupted access of the Python codes to Dropbox for reading and creation of the.txt files.

## Supporting Information

Hardware assembly, characterization of components and cells, control experiments, plasmids and bioparts, and relay switch test

## Author Contributions

SB conceived, conceptualised and designed the study. ASP, AP, and SB designed the IoT. ASP configured and assembled the IoT nodes. ASP and AP tested the IoT nodes. ASP constructed all the genetic systems. ASP, RB, BM, AP performed all the experiments. ASP and SB analysed and interpret the data and wrote the paper.

## Notes

The authors declare no competing financial interests.

## ACKNOWLEDGMENTS

This work was financially supported by SINP, Department of Atomic Energy, Govt. of India. For the JUSL deep subsurface experiments, we received assistance and inspiration from many people. We acknowledge Prof. Mala Das (in-charge, JUSL, SINP), Mr. Arindam Das (SINP), Mr. Abhishek Sen (Health Physics, BARC), Mr. Rupam Thakur (SINP), Mr. N. V. V. S Babu (UCIL) for their technical help. We acknowledge Prof. Ajit K. Mohanty (Chairman, DAE), Prof. Amit Ghosh (Director, SINP), Prof. Gautam Bhattacharya (Ex-Director, SINP), and Prof. Nabakumar Mandal (SINP) for their immense help and inspiration in initiating the biology project at JUSL.

---

**Figure S1.**
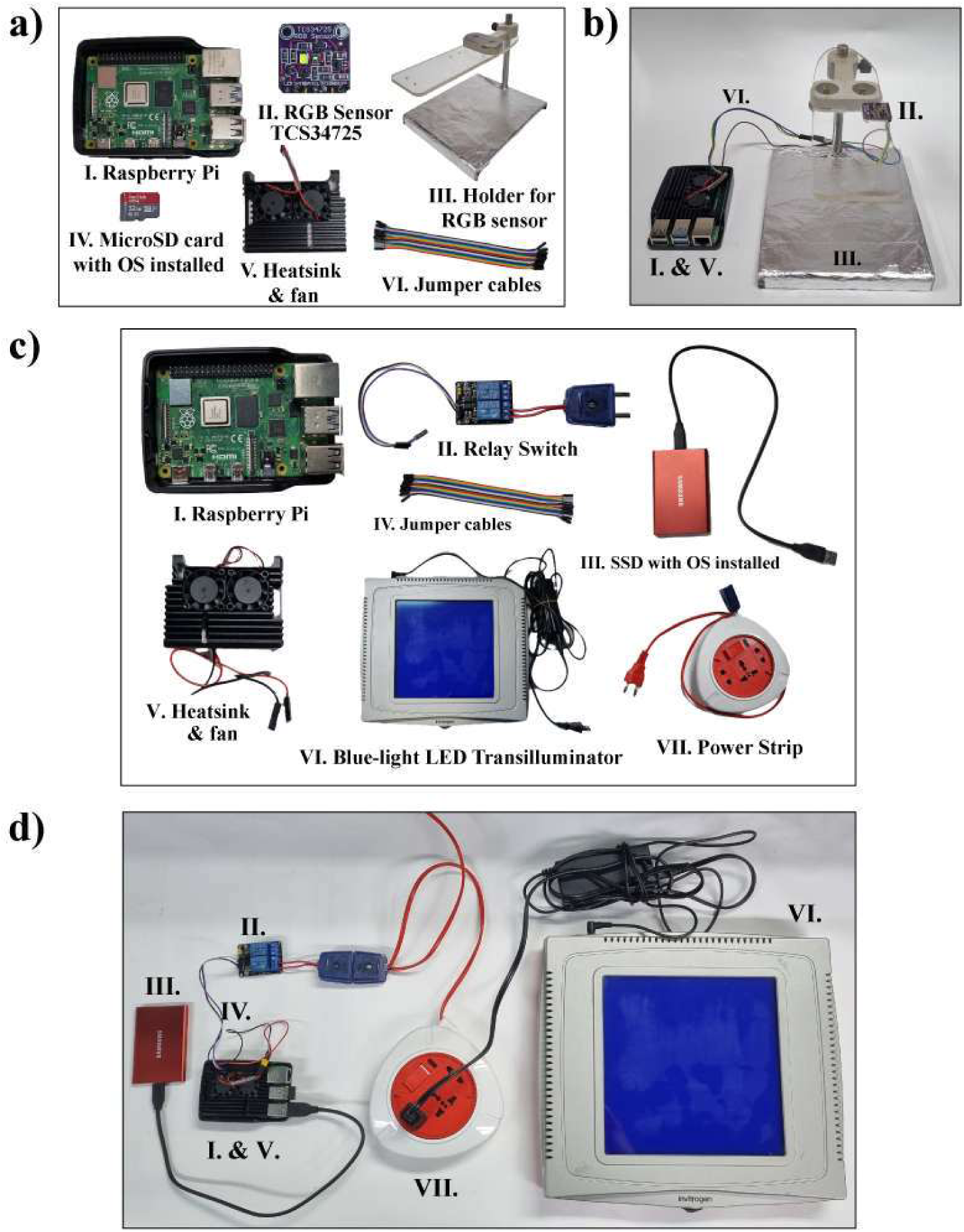
Hardware components used for construction of the IoT nodes. Accessory components like display monitors, keyboard and mouse have also been used in both the nodes. a) Individual components used in the assembly of Node Xi. b) Fully assembled Node Xi. c) Individual components used in the assembly of Node Y. d) Fully assembled Node Y.

**Figure S2.**
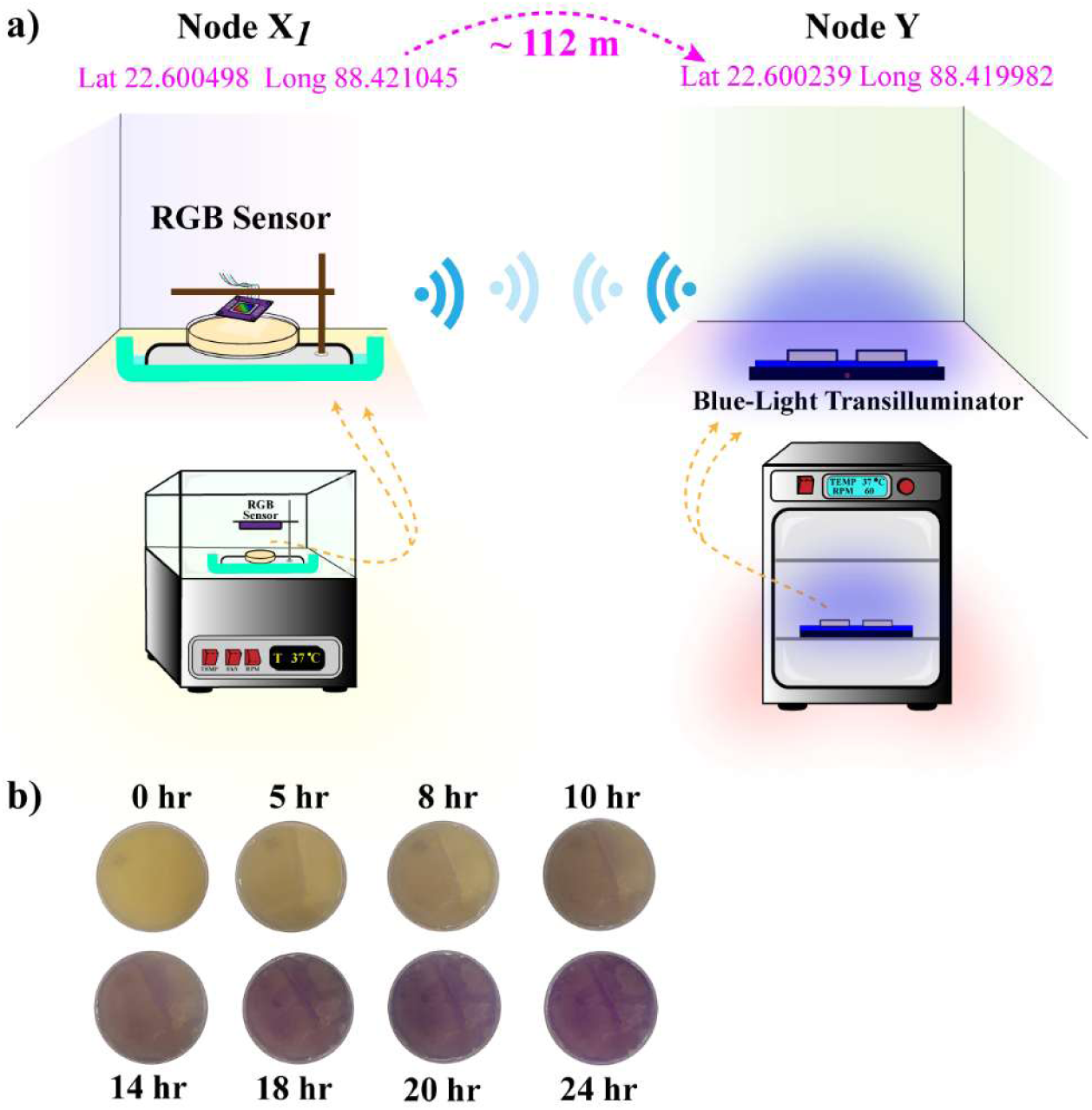
Experimental setup: Long-distance intercellular Buffer Gate. a) Schematic diagram of the experimental setup. b) A representative E2 crimson expression profile from a lawn of L-arabinose induced bacterial population as a function of time.

**Figure S3.**
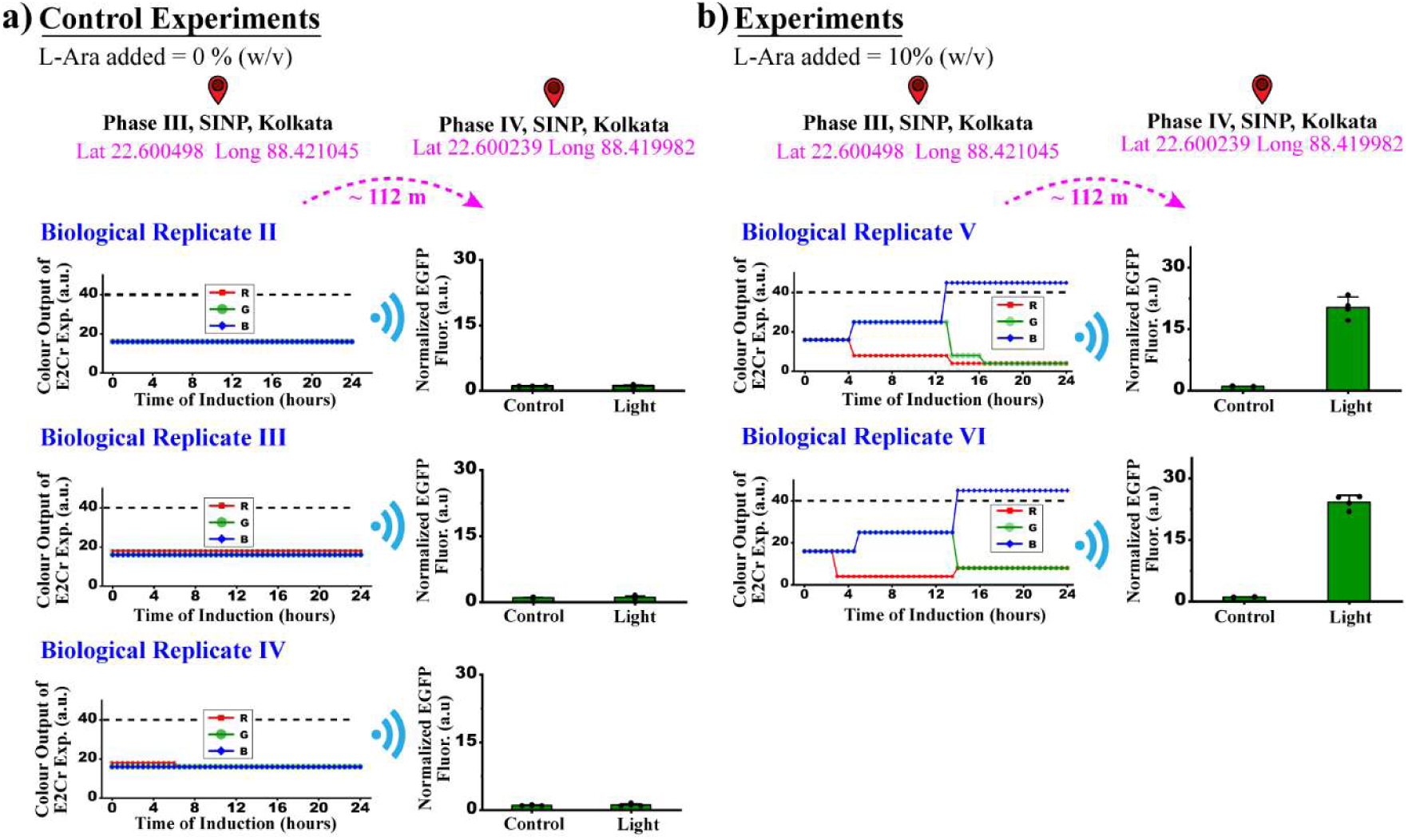
Long-distance (112 meters) intercellular Buffer gate experiments performed with with L-Ara inducible genetic circuit. a) Control experiments, no L-Ara was added to the lawn at location X1 which resulted in the absence of EGFP expression at location Y. b) Experiments where EGFP expression occurred at location Y due to E2 Crimson expression resulting from addition of L-Ara to the lawn at location X1. Apart from the one shown in figure 2., results for the two other biological replicates are shown here.

**Figure S4.**
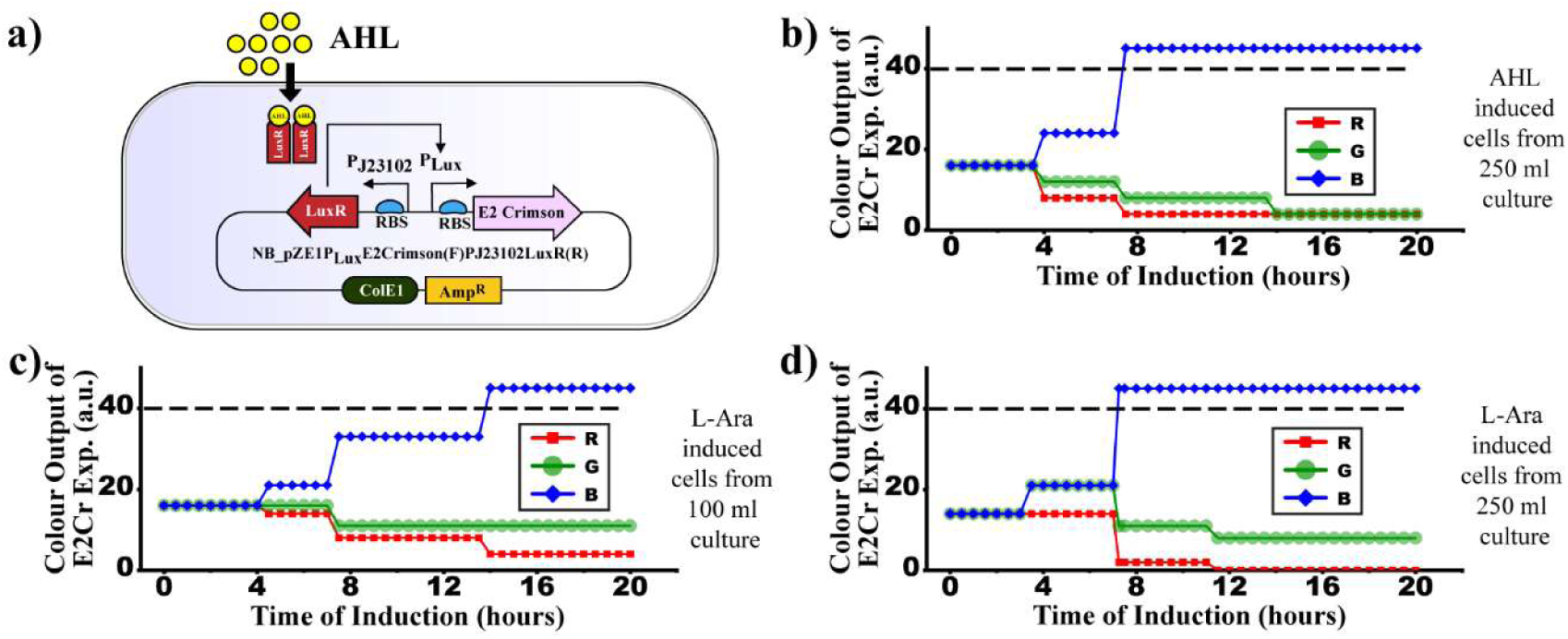
AHL-inducible genetic system and comparative gene expression colour profiles derived from lawns prepared using different numbers of bacteria. A) AHL inducible genetic system. E_2_ Crimson reporter gene kept under the control of P_Lux_ promoter in the forward direction (denoted as F) and LuxR regulatory gene has been kept under a constitutive synthetic promoter P_J23102_ in the reverse direction. b) RGB plot to illustrate lawn characterization profile when the lawn was prepared from all the AHL-inducible cells from culture volume of 250 mL. Threshold crossing was observed around 8 hours. c) RGB profile for lawn prepared with all the cells from 100 mL culture volume of L-Ara inducible cells. Threshold crossing was observed near 14^th^ hour of induction. d) RGB profile for lawn prepared from all the L-Ara inducible cells from 250 mL culture volume. Threshold crossing was observed between 7^th^-8^th^ hour of induction.

**Figure S5.**
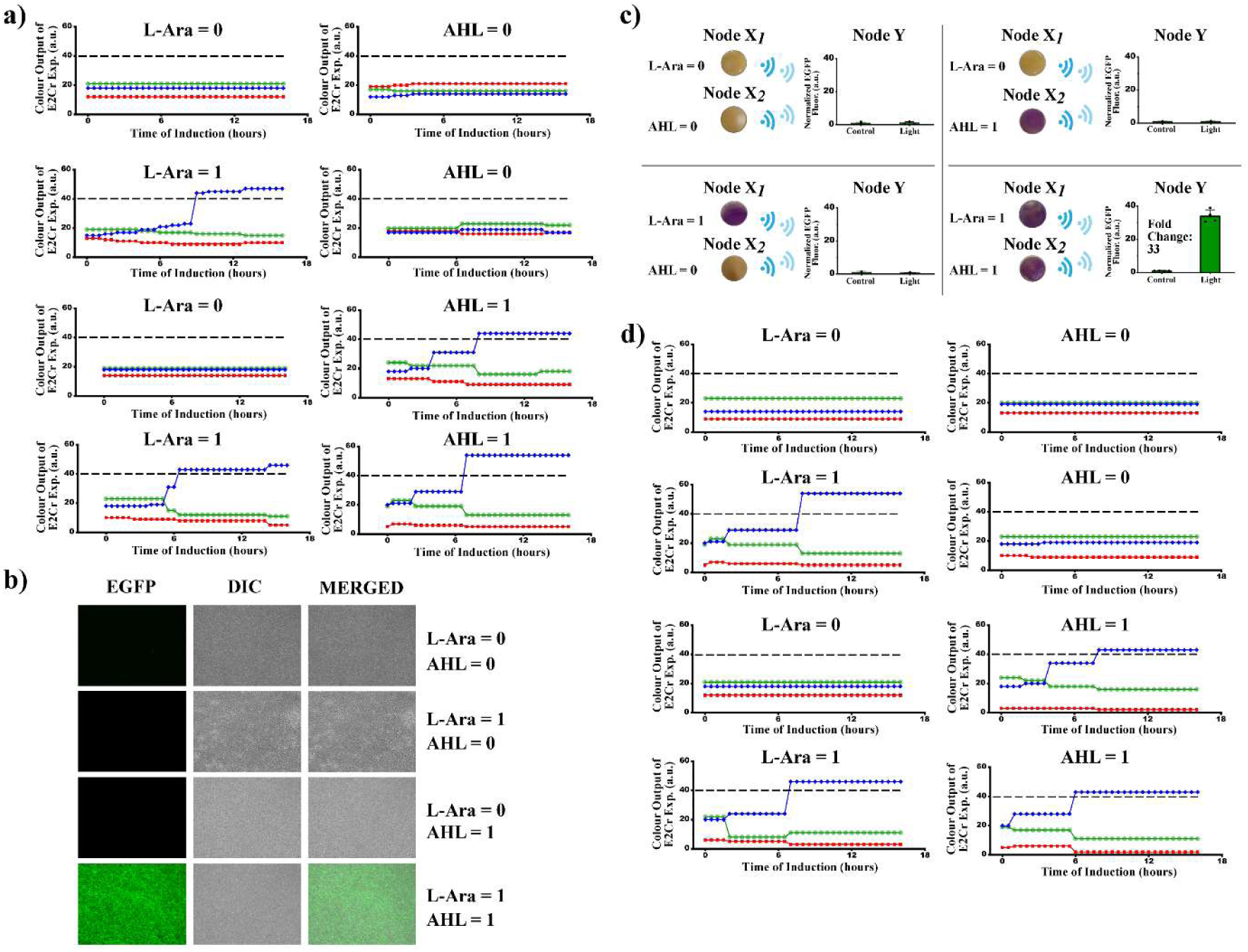
Long distance intercellular AND gate. a) RGB profile of L-Ara and AHL inducible bacterial lawn used for experiments shown in Fig. 4. b) Fluorescence (EGFP), DIC and their merged images of the fluorescence images used in Fig. 4. c) Replicates of AND gate experiment and their results. d) RGB profile for the replicates of AND gate experiments.

**Figure S6.**
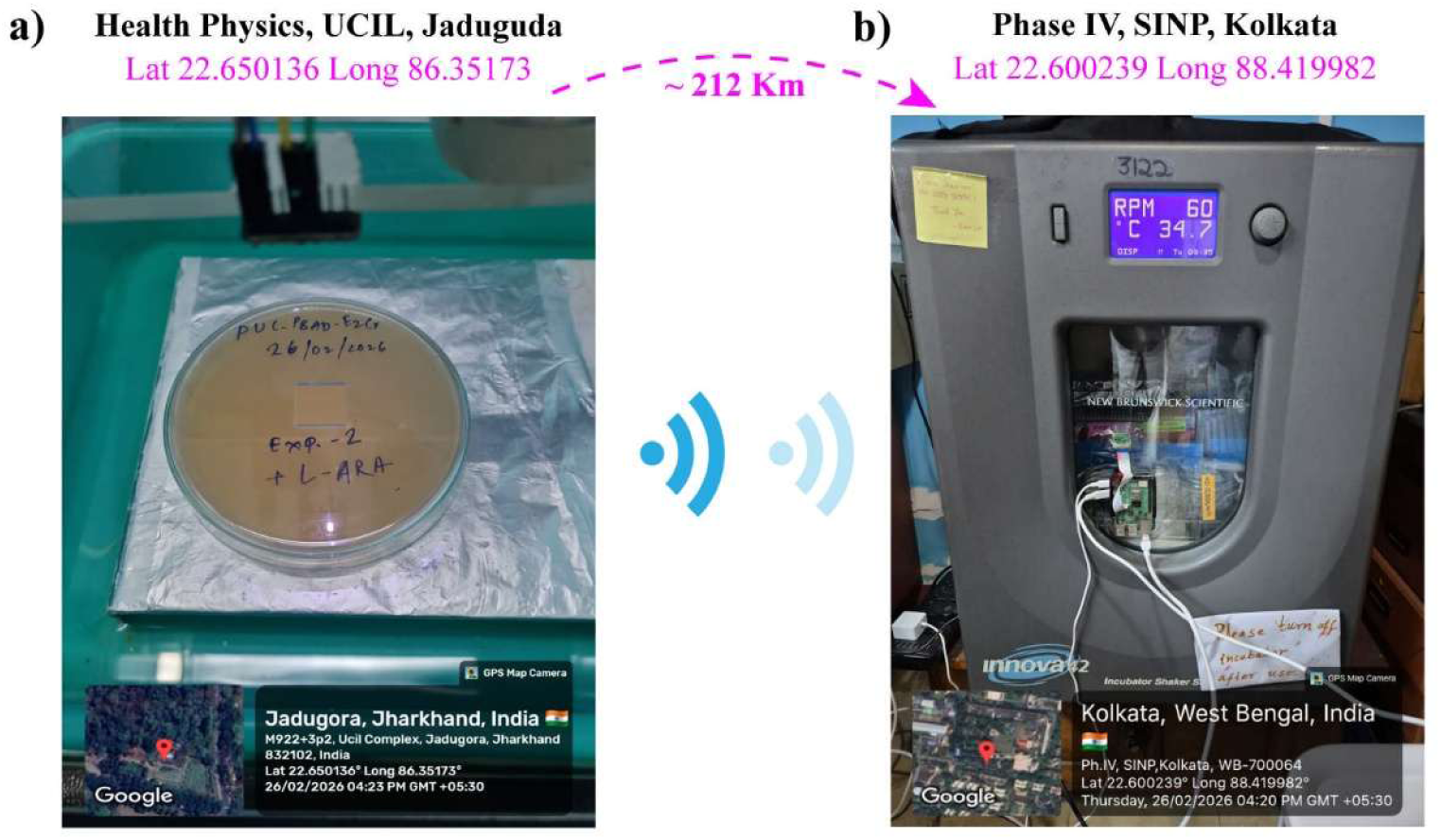
Setup for 212 km intercellular long-distance gene regulation experiments between Health Physics Surface Lab, Jaduguda and Surface Lab SINP, Kolkata. GPS information with date-time stamp can be seen in the bottom of the images. The images have been taken with an android-based smartphone camera application called “GPS Map Camera”. a) Node X1 setup at Location X1 (UCIL, Jaduguda). L-Ara is added to the lawn prepared from L-arabinose inducible cells. The lawn is kept on the platform under the RGB sensor for constant recording throughout the experiment. b) Node Y setup at Location Y (SINP, Kolkata). Cultures prepared with blue-light inducible cells on LED-light transilluminator inside an incubator. An external camera (PiCamera) can be seen mounted on the glass panel of the incubator door to take snapshots to monitor the time of blue-light switching ON.

**Figure S7.**
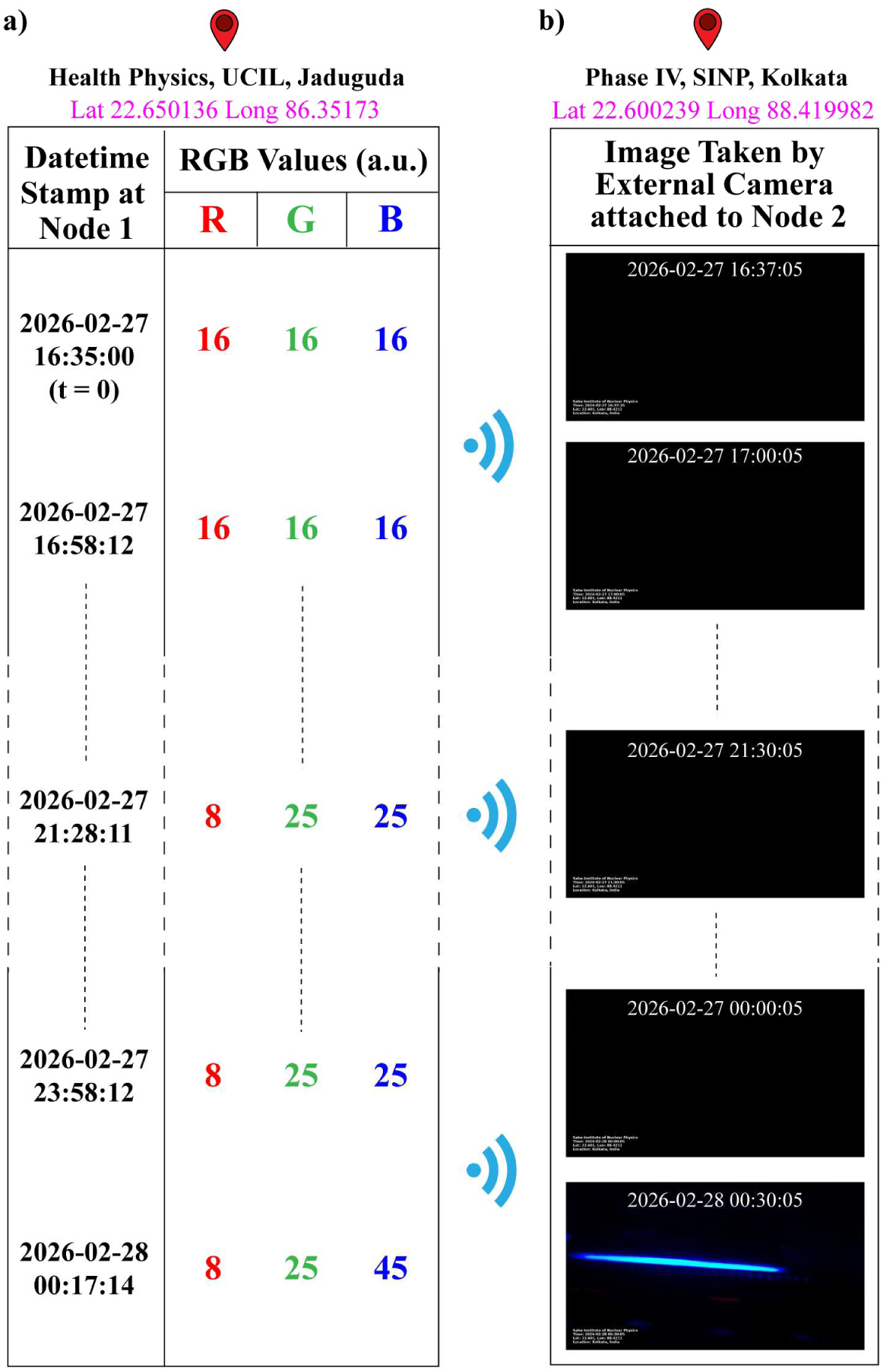
A few snapshots from 212 km experiment. RGB profile of the lawn post induction at location X1 taken at various time points and the corresponding state of blue-light (ON or OFF) at location Y at a similar timepoint. **a)** RGB values were collected from the lawn every 30 minutes after addition of the chemical inducer. The RGB values at different points are mentioned here. t_0_ indicates the time when inducer was added. ∼ 07h 40min (at 00:17:14, 2026-02-28) after addition of the inducer, B = 45 (threshold value of B>40 used throughout the study) can be seen indicating desired level of E_2_ Crimson expression at this time point. **b**) Images taken by an external camera attached to Node Y at location Y. The blue light can be seen in its OFF-state (first four images) as long as B < 40 at location 1. When desired E2 Crimson expression is detected by the RGB sensor, the blue light is switched ON (5^th^ image in the panel). The camera was programmed to take a snapshot with date-time stamp and GPS information every 30 minute.

**Figure S8.**
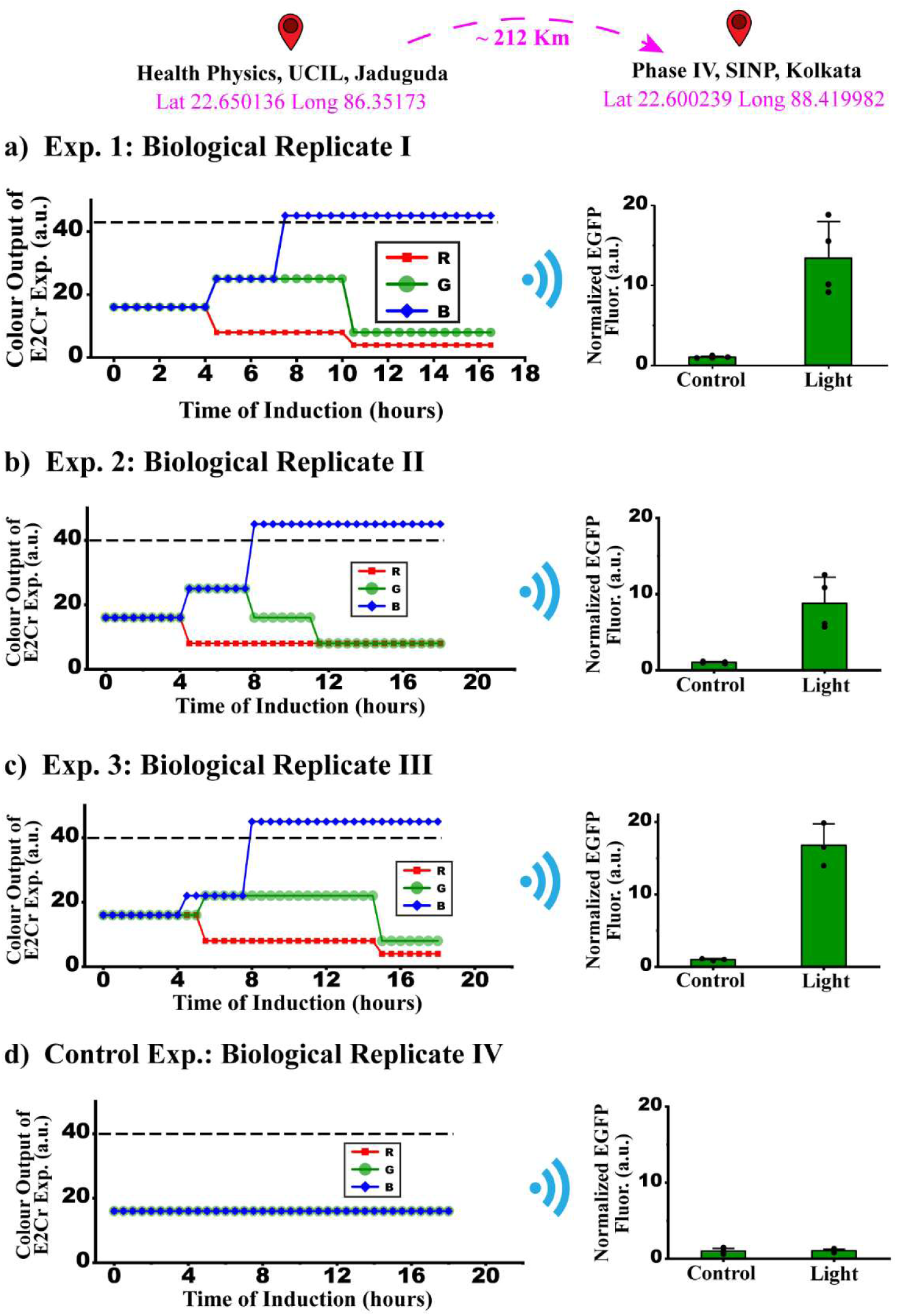
Testing the distance scalability of IoGT system between two cities located 212 Km apart. **a)** A set of experiment where L-Ara was added to the lawn at the first location. Corresponding EGFP expression observed at location Y. b) Second biological replicate where L-Ara was added. c) Third biological replicate with L-Ara. d) Control experiment where no L-Ara was added to the lawn. No EGFP expression observed at location Y.

**Figure S9.**
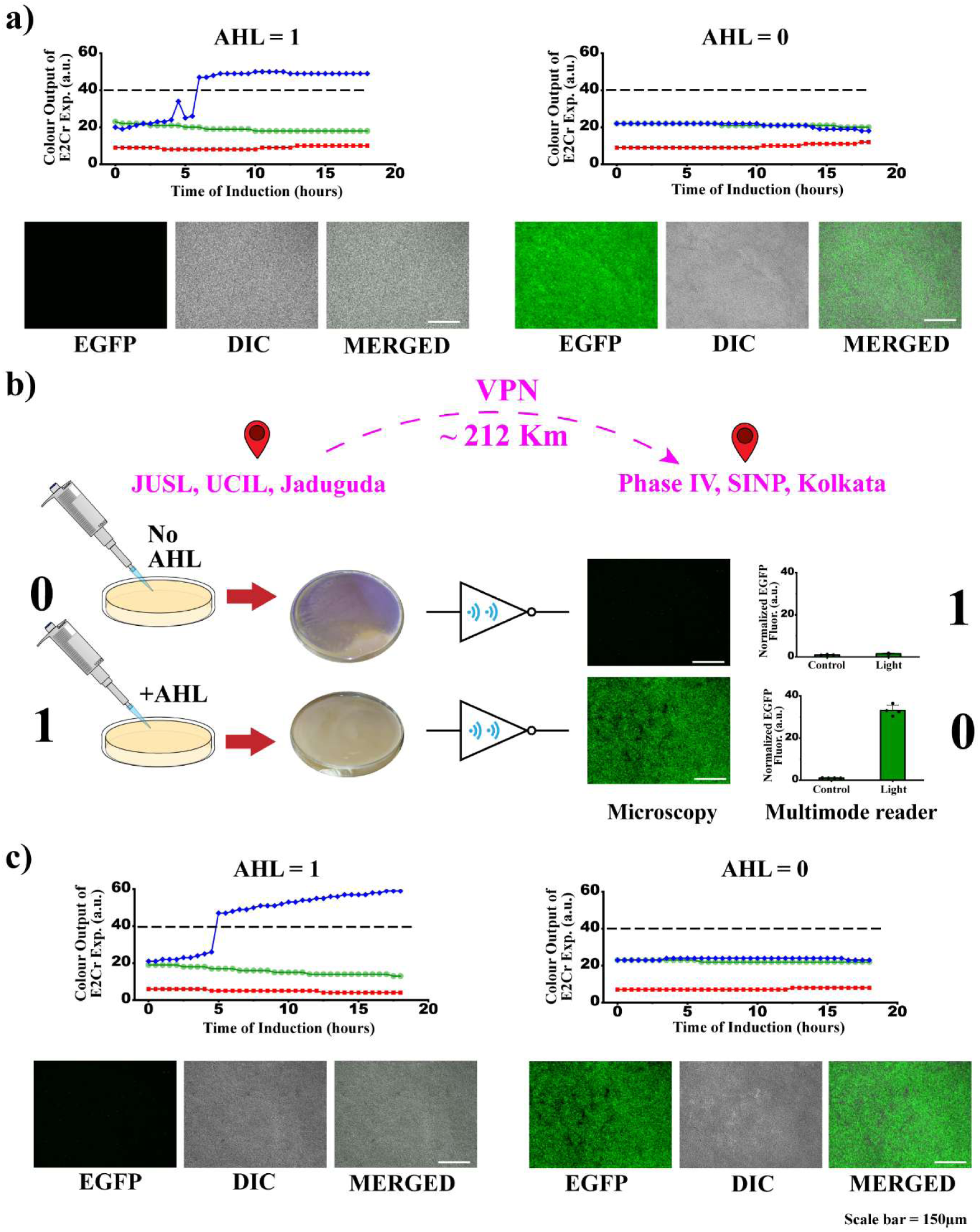
Long distance intercellular NOT gate. a) RGB profile of the bacterial lawns and fluorescence, DIC and their merged images of the microscopy result shown in Fig. 5. b) Replicate of long distance NOT gate. c) RGB profile of the bacterial lawns and fluorescence, DIC and their merged images of the microscopy result for the replicate of long distance NOT gate.

**Figure S10.**
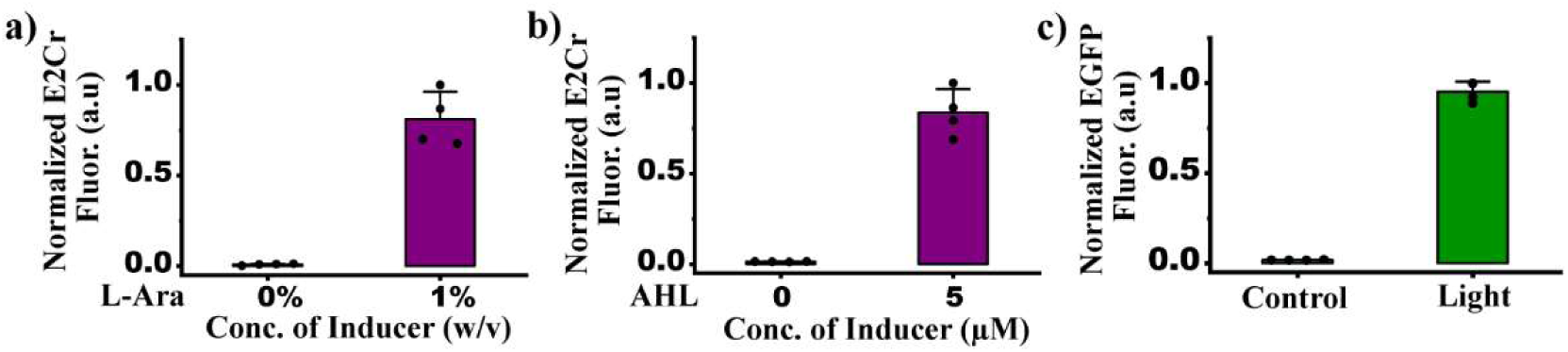
**Initial experimental characterization of the genetic circuits used in the study in liquid culture**. E2 Crimson is denoted as E2Cr in the plots. a) L-Ara inducible genetic circuit was initially characterized in liquid LB media with appropriate antibiotic and 1% L-Arabinose for induction. Fluorescence measurements were taken 16 hours post induction. b) AHL – inducible genetic circuit was also initially characterized in liquid LB media with appropriate antibiotic and 5 µM C6-HSL for induction. Fluorescence measurements were taken after 16 hours of induction. c) Initial characterization of pDawn-EGFP circuit was performed by 100 times dilution of overnight culture (in LB, Miller) of cells transformed with the circuit in M9 minimal media and blue-light induction for 6 hours.

**Table S1.** RGB values as a function of sensor distance.

| Distance between Sensor and object (mm) | RGB Values (a.u.) |  |  |
|---|---|---|---|
|  | For Red Paper |  |  |
|  | R | G | B |
| 0 | 0 | 0 | 0 |
| 2 | 30 | 11 | 11 |
| 5 | 45 | 8 | 8 |
| 7 | 92 | 16 | 16 |
| 10 | 255 | 0 | 0 |
| 11 | 255 | 0 | 0 |
| 12 | 255 | 0 | 0 |
| 15 | 0 | 0 | 0 |
| 19 | 0 | 0 | 0 |
| 25 | 0 | 0 | 0 |
|  | For Green Paper |  |  |
|  | R | G | B |
| 0 | 0 | 0 | 0 |
| 2 | 2 | 28 | 16 |
| 5 | 4 | 40 | 10 |
| 7 | 11 | 30 | 11 |
| 9 | 68 | 45 | 6 |
| 10 | 0 | 45 | 8 |
| 12 | 0 | 45 | 0 |
| 15 | 0 | 0 | 0 |
| 20 | 0 | 0 | 0 |
| 25 | 0 | 0 | 0 |
|  | For Blue Paper |  |  |
|  | R | G | B |
| 0 | 0 | 0 | 0 |
| 2 | 0 | 0 | 45 |
| 5 | 0 | 8 | 45 |
| 7 | 0 | 0 | 45 |
| 10 | 0 | 0 | 255 |
| 11 | 0 | 16 | 92 |
| 13 | 0 | 0 | 0 |
| 15 | 0 | 0 | 0 |
| 20 | 0 | 0 | 0 |
| 25 | 0 | 0 | 0 |

**Table S2.** Description of plasmids constructed for this study.

| **SL. No.** | **Plasmid** | **Description** | **Cassette(s)** | **Origin** | **Anti-biotic Select-ion** | **Purpose** | **Source** |
|---|---|---|---|---|---|---|---|
| **1.** | NB_pZU1_P<sub>BAD</sub>_R1_E2_Crimson | L-Ara inducible plasmid | P<sub>BAD</sub>-RBS1-E2_Crimson-Terminator | pUC | Amp | Long-Distance Commun-ication | This study |
| **2.** | NB_pZE1P<sub>Lux</sub>_R1_E2_Crimson(F) J23102_R1_LuxR(R) | AHL inducible plasmid | P<sub>Lux</sub>-RBS1-E2_Crimson-Terminator (Forward)<br>P<sub>J23102</sub>-RBS1-LuxR-Terminator (Reverse) | ColE1 | Amp |  |  |
| **3.** | pDawn-EGFP | Light inducible plasmid | EGFP | pUC | Kan |  |  |

**Table S3.** Bioparts and their source to build the plasmids used in this study.

| **SL. No.** | **Plasmid** | **Assembly** | **Purpose** | **Source** |
|---|---|---|---|---|
| **1.** | pNB_EA_3EGFP | Network Brick | To get NB vector with pUC origin and EGFP reporter gene | Mukhopadhyay et al., 2020 |
| **2.** | pM2_P<sub>BAD</sub>_R1_EGFP |  | To get NB vector backbone with ColE1 origin and P<sub>BAD</sub> promoter | Bonnerjee et al., 2024 |
| **3.** | pK1_PLux_R1_cI* |  | Source of P<sub>Lux</sub> promoter |  |
| **4.** | pM2_J23102_R1_LuxR |  | Source of the cassette J23102-RBS1-LuxR-Terminator |  |
| **5.** | pK1_PR_R1_E2 Crimson |  | Source of E2 Crimson reporter gene |  |

**Table S4.**
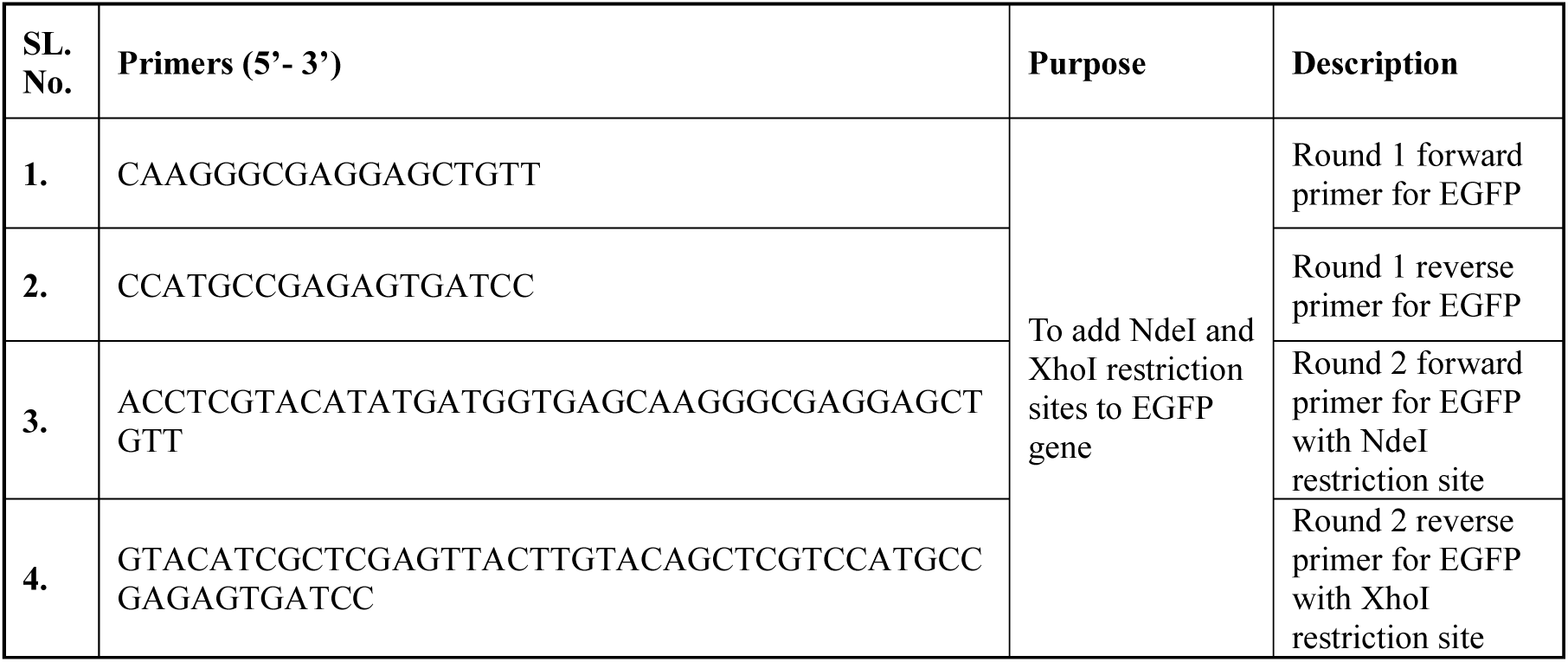
Details of oligos used in this study.

